# Representations of spatial saliency in auditory cortex are selectively organized through temporal coordination

**DOI:** 10.64898/2026.06.10.731488

**Authors:** Martin Irani-Cereceda, Zhili Qu, Kamal Sen, Sepideh Sadaghiani, Howard J. Gritton

## Abstract

Selective processing of behaviorally relevant sounds in complex environments requires dynamic modulation of auditory representations. Here, we show that behaviorally relevant sound locations selectively organize auditory cortical activity through temporal coordination rather than changes in firing rate. Using laminar electrophysiology in mice, we recorded activity in primary auditory cortex (A1) activity during a task in which reward was associated with a specific sound location while neutral, non-rewarded sounds were presented across multiple spatial positions. Neurons exhibited increased response reliability, temporal precision, and spike-field coupling to gamma oscillations selectively for sounds presented from behaviorally relevant locations, even when those sounds were unrewarded. These effects were absent during passive listening. Neurons with stronger gamma coupling also had sharper spatial tuning and greater trial-to-trial reliability, linking temporal coordination to representational fidelity. These findings identify gamma-mediated synchronization as a mechanism through which spatial relevance dynamically reshapes auditory encoding.

## INTRODUCTION

Sound localization is fundamental to effective auditory scene analysis, enabling listeners to segregate competing sources based on spatial location^11,41^. In the primary auditory cortex (A1), most neurons are sensitive to sound location, typically exhibiting broad receptive fields and stronger contralateral tuning^53^. Rather than relying on sharply tuned, topographically organized maps, spatial information however, is represented through distributed ensemble activity across auditory cortical populations^84^. This distributed organization creates a fundamental computational challenge for the auditory system. In contrast to visual cortex, where spatial attention can selectively amplify neuronal population firing rates corresponding to specific retinotopic coordinates^76^, the broadly tuned and heavily overlapping auditory spatial representations are difficult to amplify through simple gain amplification. How then does auditory cortex dynamically prioritize behaviorally relevant regions of space? Resolving this question is critical for understanding how listeners selectively extract relevant sound source information in complex acoustic environments during auditory scene analysis.

A1 populations rely on two complementary coding strategies to represent acoustic information^2,80^. For slow time-varying signals, synchronized neuronal populations encode stimulus features through precise spike-timing, with action potentials phase-locked to the temporal stimulus modulations^80^. Beyond individual firing rates, neuronal populations can coordinate the relative timing of their spikes to enhance relevant stimulus features^2,20,63^. This spike-timing coordination carries information about acoustic features that cannot be recovered from firing rate alone^20,39^. Although temporal coding has traditionally been studied in the context of acoustic feature representation, recent evidence suggests that it also contributes substantially to spatial encoding, carrying information about the spatial origin of sounds^59,69^. These observations suggest that spatial information encoded subcortically by rate may alternatively be carried within cortical population through temporal dynamics rather than rate coding alone at the level of the cortex^24,59^. However, most of our present understanding of auditory cortical spatial processing comes from studies of rate coding features^17,45,75,82^, leaving unresolved whether behaviorally relevant spatial information is primarily represented through temporal coordination, firing-rate modulation, or both. Importantly, if rate alone cannot account for selective spatial processing in distributed auditory representations, then temporal coordination may provide an important alternative coding dimension through which behaviorally relevant locations can be prioritized.

A candidate mechanism for such temporal coordination is oscillatory synchronization. Oscillatory fluctuations in extracellular potentials reflect alternating periods of excitation and inhibition within neuronal populations and have been proposed to organize neural dynamics by aligning neuronal excitability, routing information between cortical circuits and coordinating cognitive control processes^14,28,55^. Oscillatory entrainment to stimulus structure and behavioral rhythms further supports a role for temporal coordination in auditory processing^15,43,44^. In particular, local gamma-band oscillations (40-60Hz) can enhance network selectivity to feedforward inputs and regulate the temporal precision of neuronal responses^46,61^. Oscillatory coordination is especially attractive as a mechanism of auditory spatial selection because it permits dynamic routing of distributed neuronal ensembles without requiring spatially arranged maps. Rather than amplifying fixed spatial channels, temporal synchronization may allow behaviorally relevant representations to emerge transiently from distributed cortical populations. Within this framework, gamma synchronization could facilitate temporal binding in a way that transiently links encoding of behaviorally relevant locations into coherent functional ensembles at the neuronal level.

These observations taken together provide support for the possibility that relevant behavioral locations could be spatially modulated through temporal coordination in the auditory cortex. Such processes could also function independently from firing rate modulation. Here, we tested this hypothesis by examining whether behaviorally-relevant sound locations modulate auditory cortical representations through changes in temporal coordination at both the population and single unit level. Using laminar recordings spanning multiple cortical depths in behaving mice, we measured neural and local field potential (LFP) activity during a task in which reward was consistently associated with a specific spatial location. Because A1 neurons also carry reward and prediction signals^12^, we probed cortical activity using neutral, non-rewarded amplitude-modulated (AM) sounds presented randomly from multiple spatial locations throughout the task. This approach allowed us to isolate sensory processing from reward- and motor-related activity. We found that behaviorally relevant locations selectively enhanced gamma synchronization, spike–field coupling, response reliability, and spatial tuning independent of consistent firing-rate modulation. Together, these findings support a model in which temporal coordination dynamically organizes auditory spatial representations according to behavioral relevance and identify gamma-mediated synchronization as a candidate circuit mechanism for flexible auditory scene analysis.

## RESULTS

### Behavioral engagement establishes spatially selective saliency

To study how representations of sound sources in A1 are modulated by location relevance, we trained water restricted head-fixed mice to lick a water port in response to a 10kHz pure tone to receive a reward (**Fig. 1A**). Once well trained, animals underwent electrophysiology recording in an anechoic chamber on a sound localization task using four speakers arranged along the azimuth (0°elevation relative to the animal but spanning 180°along the azimuth: see Methods; **Fig. 1B**) In addition to delivery of the rewarded pure tone stimulus, mice received neutral probe sounds consisting of amplitude-modulated (AM) broad band noise shaped by the envelope of speech tokens randomly from the four speaker locations. The use of speech tokens preserved the underlying temporal structure inherent in communication (i.e., prose, syllable, and phonetic structure) but convolved with white noise to enhance recruitment of A1 populations in mice (**Fig. 1C-1D**). Although the AM envelopes were derived from human speech, the spectro-temporal features encoded by auditory cortical neurons operated on timescale consistent with those of mouse ultrasonic vocalizations, supporting the relevance of these stimuli for probing auditory cortical processing^59^.

**Figure 1.**
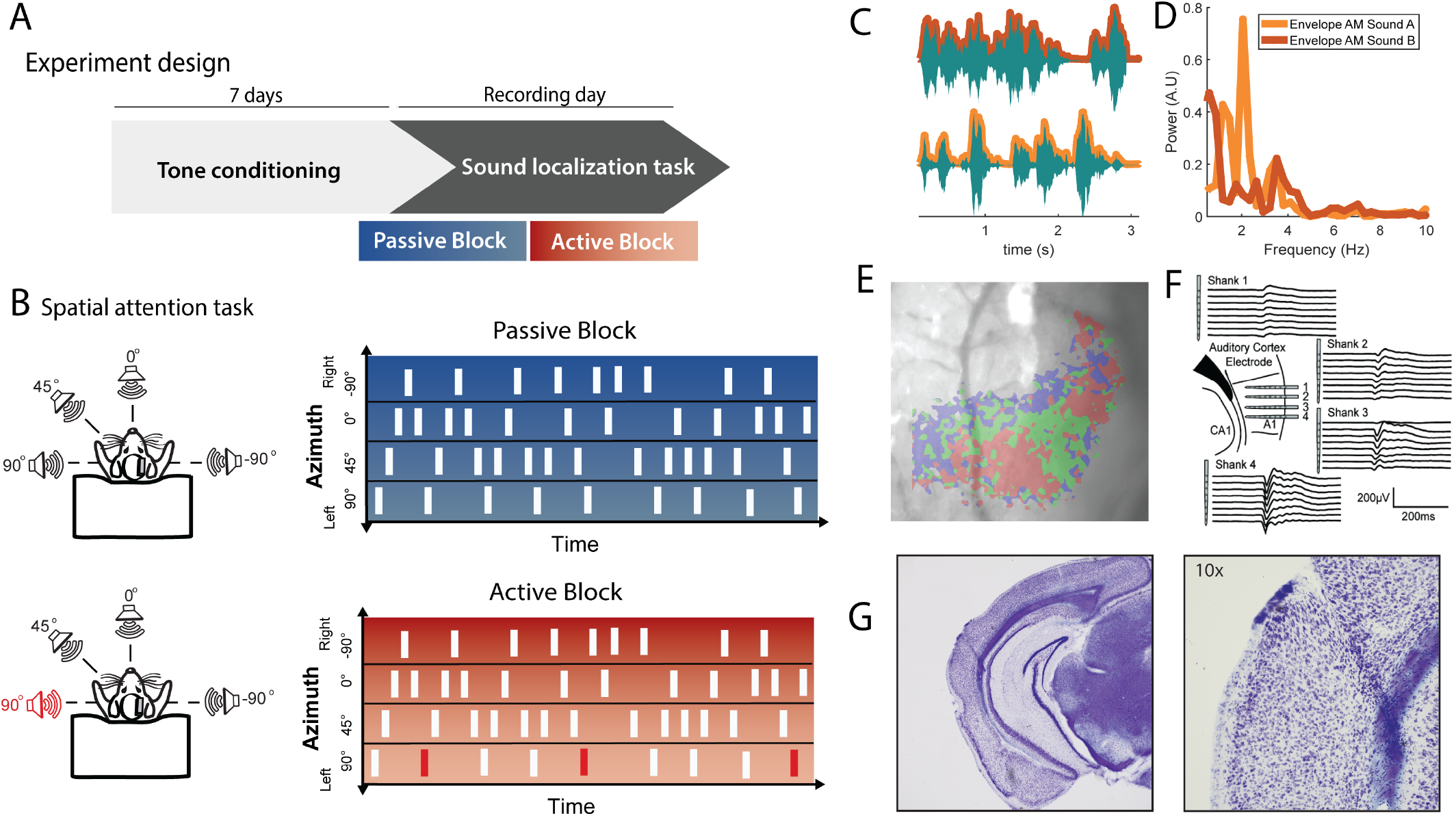
Experimental Design. **A**. Training protocol. We trained mice to conditionally lick to obtain a small water reward in response to a 10 kHz tone. Training occurred over 7-10 continuous training days before animals were recorded in the spatial localization task. All mice reached asymptotic performance of 162 rewards and no misses before moving to the recording stage of the task. **B**. After training, animals moved to a task consisting of a block design where animals would be presented with both the rewarded tone and two AM modulated white noise (sound A or sound B) coming from any of four speakers arranged along the azimuth. A passive block would only consist of AM sounds, whereas active blocks would contain the 10 kHz rewarded tone presented occasionally but only at the rewarded location (90 degrees). **C**. Temporal traces showing the two AM sounds that were used in this study with their envelopes outlined. **D**. Power spectral density of sound envelopes for the two sounds shown in (C). **E**. Intrinsic signal optical image of A1 from an example mouse used in this study under anesthesia and prior to task training. Shaded areas delineate sound modulated areas for targeting of electrode to A1 using pure tone frequencies. Maximal responses for each of the different tone frequencies (red: 3 kHz; green: 10 kHz; blue 20 kHz) are shown. **F**. Mice were implanted with a 4-shank, 32-channel electrode array, in the right hemisphere of A1 area based on intrinsic imaging maps. Each shank contained 8 sites per shank with 100 µm spacing between electrode contacts. **G**. Representative image from a Nissl-stained histological section showing electrode placement in A1 confirmed following histological processing.

During the task, mice experienced playback of sounds within two contexts: in a passive listening context, where only neutral sounds occurred, and an active listening context where between every two to five neutral sound trials, a pure tone originating from one sound source location (rewarded location) was played. Active and passive sessions were presented in counterbalanced order (before or after) across sessions and animals, and the combined data is presented in the results. During the active listening context described in this experiment, the pure tone sound was presented at the 90°location to take advantage of the fact that the contralateral field would be more strongly represented in the right hemisphere^53^ of A1 where electrodes were placed in all mice (**Fig. 1E-G**). A1 electrode placement was determined by intrinsic signal imaging prior to implantation and confirmed with histology at the conclusion of the study (**Fig. 1E, Fig. S1**; see methods). We reasoned that animals would selectively attend towards salient locations that were paired with pure tone-reward delivery, and that this effect would be reflected in both the representations for the neutral stimuli as well as the rewarded stimuli. Consequently, we could covertly prime spatial selectivity by comparing processing of the neutral AM sound across contexts (passive and active) and location (rewarded and non-rewarded). The analysis of neutral stimuli within our experimental design also allowed us to isolate spatial modulation from reward and motor (licking) confounds.

We first characterized behavioral responses, spectral power, and pupil dynamics across passive and task-engaged conditions. To probe task engagement, we evaluated stimulus and location specific responding (hit rates) to identify whether in the task-engaged conditions, mice selectively responded only for the rewarded tone, and that responding was largely absent for the AM sound presentations regardless of the location it was played (**Fig. 2A-2B**). We found that mice learned to generally refrain from licking for the AM stimulus in the active block and in the rare instances of licking, responses did not differ across the four speaker locations (one-way ANOVA; *F*_(3,64)_ = 0.077, p = 0.972). In addition, we found that animals responded 93.25±0.08% (mean + std. dev.) to tones presented during the active sessions suggesting a highly maintained level of engagement for each recording session (**Fig. 2C**). Notably, this engagement was also associated with an enhancement in alpha/beta (8-20Hz) power during the inter-trial interval (ITI) in the active session compared to the passive condition (**Fig. 2D**; two-sided sign-rank Wilcoxon test, z = 2.9586, p = 0.0031). Elevated alpha power is associated with task engagement and contributes to prioritizing relevant information^68^. Because pupil diameter can be used as a proxy associated with changes in arousal, listening effort, and reward magnitude^34,83^, we quantified pupil changes in response to the neutral stimulus sound presentation across active and passive contexts for a subset of mice in the experiment (n=4; **Fig. 2E-H**). Pupil diameter was quantified using the area under the curve (AUC) from images triggered by the electrophysiological equipment and collected during sound playback for the AM neutral sounds across the four locations. A two-way ANOVA including subject as a random factor examined the effects of location and condition on pupil diameter. Analysis revealed a significant main effect of location, *F*_(3,9)_=5.33, *p* = 0.022, but no significant effect of condition, *F*_(1,9)_=2.09, *p* = 0.244. Importantly, there was a significant interaction between location and condition, *F*_(3,9)_ = 4.98, *p*=0.026, indicating that the effect of location on pupil diameter differed across passive and task-engaged states. The dissociation between tonic alpha-beta enhancement during the ITI and phasic-location specific pupil responses suggest that behavioral engagement during the active block operates at two complementary processes where tonic engagement involves preparatory cortical states and a phasic arousal response captured by pupil diameter that transiently occurs during salient spatial inputs.

**Figure 2.**
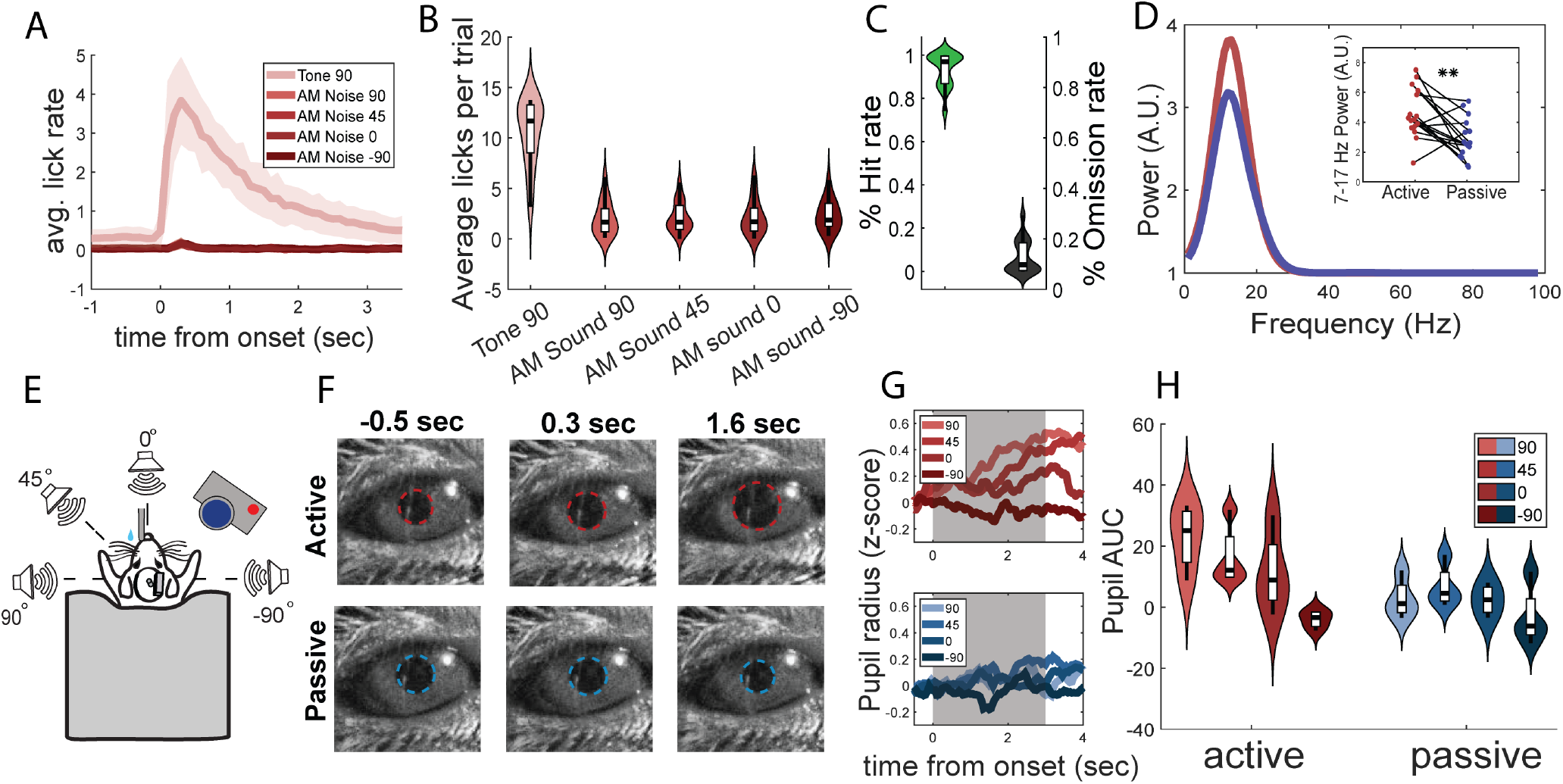
Reward availability increases behavioral saliency of sound source location. **A**. Average lick rate traces in response to tone presentation and AM sound presentations across all four locations. **B**. Average lick rate during AM sound presentations in active blocks separated by stimulus and by location. Note elevated licking to the 10kHz tone that is not present for the AM sounds. **C**. Animal hit and omission rates for the 10 kHz tone presentations from each recording session. **D**. Power spectral density of pre-stimulus activity in active and passive blocks differs across behavioral states. Inset: Power in alpha-beta frequencies (7-17 Hz) observed in baseline (previous two seconds before sound onset) during active and passive block. We observed a significantly higher alpha-beta power in active with respect to the passive state (** indicate *p <* 0.001). **E**. Diagram of experimental set up including speaker location, camera for pupil tracking, and water port. **F**. Sequential sample frames following stimulus onset recorded from a representative mouse in active (*top*) and passive (*bottom*). **G**. Average diameter during sound playback for each sound source location. H. Average AUC of pupil diameter over time for each sound source location.

### Behavioral saliency of sound source location augments gamma-band activity and theta-gamma coupling in auditory cortex

We next asked whether spatial saliency modulates sound-evoked cortical rhythms and population activity in A1. To do so we recorded local field potentials and single unit activity using 4-shank, 32 channel laminar electrode arrays spanning all cortical layers in mouse A1 (**Fig. 1E, Fig. 3A**). A cluster-based permutation test of spectral power of local field potentials revealed a significant increase in alpha-beta (15-30 Hz; cluster corrected *p <* 0.01) and gamma band power (40-60 Hz; cluster corrected *p <* 0.01) during the active behavioral block relative to the passive block (**Fig. 3B**). We further examined the power of these bands across all four speaker locations in the active block using a linear mixed-effects model with location as a fixed effect and subject as a random effect (**Fig. 3C**). We observed a significant effect of location on gamma power (*β* = -0.078, SE = 0.027, *t* = -2.95, *p* = .004, 95% CI [-0.131, -0.025]). No effect of location was observed for alpha-beta. Moreover, we found that gamma activity was markedly increased in superficial channels in A1 (**Fig. 3D**), this was supported by the current source density analyses of gamma events, where we observed localized gamma activity at intermediate recording depths propagating towards superficial channels for each shank (**Fig. S2C**). In addition, gamma burst event activity tracked the temporal envelope of amplitude-modulated stimuli, indicating that oscillatory dynamics are aligned with stimulus structure (**Fig. 3G, Fig. S2A-B**) in direct support of cortical tracking of sounds by gamma activity.

**Figure 3.**
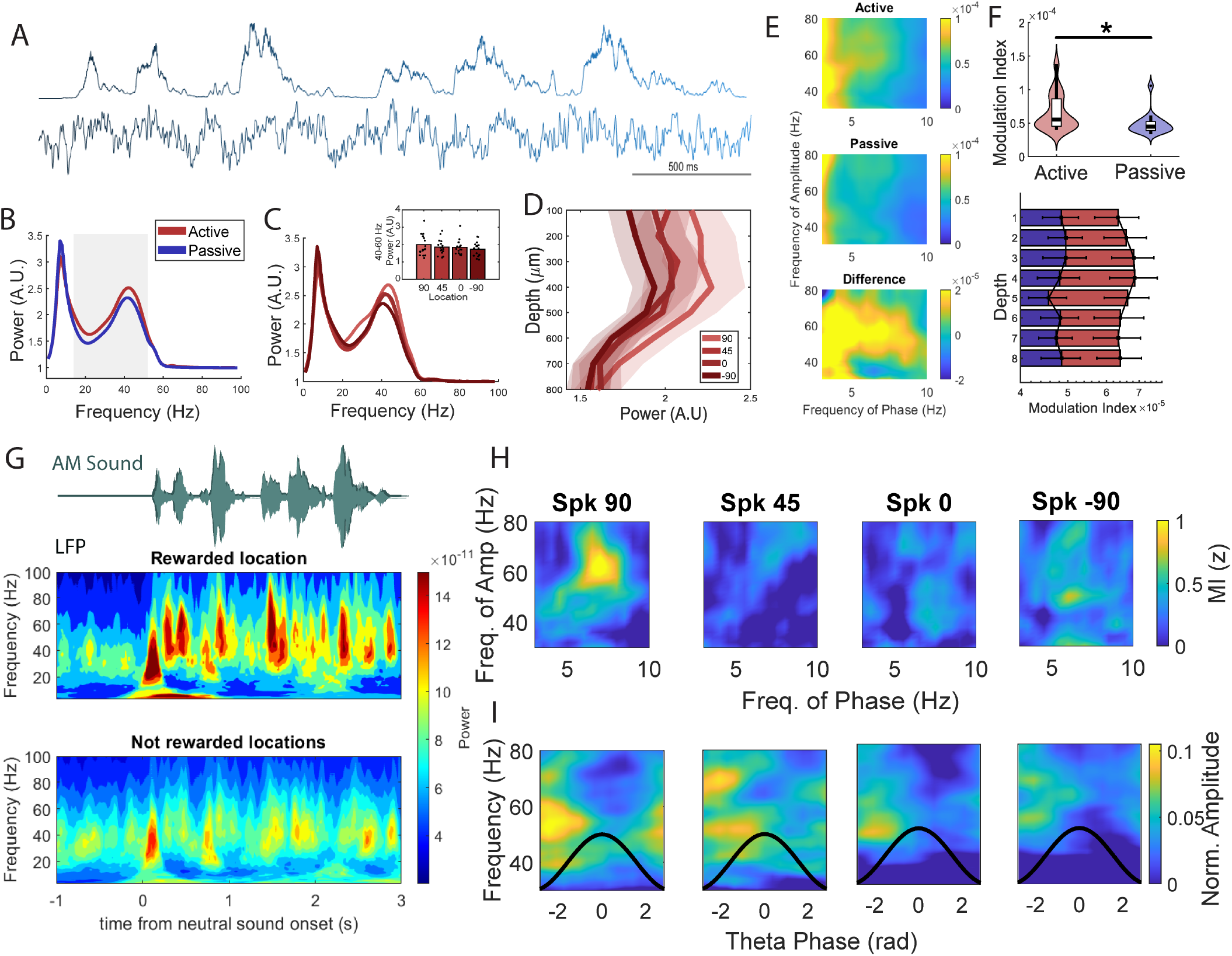
Oscillatory increases in gamma and theta-gamma coupling are state-dependent and aligned to behavioral saliency of spatial location. **A**. Envelope of one of the two AM sounds (*top*) and a representative LFP collected from a single channel (*below*). **B**. Power spectral density for active (red) versus passive (blue) conditions. Shaded area denotes significant differences between conditions. **C** Power spectral density for each location in the active period. *Inset* : bar plots showing power for 40-60 Hz frequency band. **D**. Sound evoked depth profile of gamma (40-60 Hz) power for each spatial location, where gamma was noted to be elevated across spatial locations only in superficial channels. **E**. Comodulograms with raw modulation indices for active (*top*) and passive (*middle*) conditions, and difference (*bottom*) between both conditions. **F**. Violin plots denoting average theta-gamma phase-amplitude coupling MI for each subject (*top*). **G**. Representative spectrogram for rewarded (*top*) and non-rewarded locations (*bottom*) in the active block.**H**. Z-scored modulation indices in active block for each spatial location. **I**. Cross frequency coupling between theta phase (4-8 Hz) and high frequency power during sound playback. Solid black lines denote one cycle of theta. We found that higher frequency coupling (40-70 Hz) was organized primarily to the rising phase of the theta cycle.

LFPs also showed strong theta oscillations that peaked near 7 Hz and were phase locked to slower components of the envelope for AM sounds (**Fig. 3A**). Theta power did not vary significantly between conditions or across locations (**Fig. 3B-C, Fig. S3B**). However, we observed a modulation of gamma amplitude by theta phase in the active block that was significantly higher than the passive block (**Fig. 3E-F**, two-sided Wilcoxon signed-rank test, *p* = 0.0352). We observed coupling of gamma and theta across depths (**Fig. 3F**). Using a linear mixed-effects model, relative to the 90° location (intercept estimate = 0.50 ± 0.09 SE), this cross-frequency modulation was significantly reduced at 45°(*β* = -0.36 ± 0.11, *t* = -3.37, *p* = 0.0013) and 0°(*β* = -0.31 ± 0.11, *t* = -2.84, *p* = 0.006). Modulation at the ipsilateral location was also lower (*β* = -0.15 ± 0.11, *t* = -1.41), although this difference did not reach statistical significance (*p* = 0.163; **Fig. 3H**). We analyzed the amplitude of higher frequencies with respect to the phase of theta and found that gamma had the highest magnitude during the rising theta phase and lowest during the falling phase (**Fig. 3I**). Together, these findings demonstrate that behavioral relevance selectively enhances gamma-band synchronization in auditory cortex and that gamma modulation is coupled to theta rhythms in the auditory cortex during task-engagement.

### Gamma synchronization organizes spike timing and enhances response reliability

To determine whether spatially modulated oscillatory dynamics influenced neuronal spiking, we quantified spike–field coupling across frequencies. Specifically, we used pairwise-phase consistency (PPC), which measures the strength of spike-phase coupling as a bias free and firing rate independent estimator^79^. We used the adjacent recording sites to each analyzed single neuron site to avoid spike bleed-through in the LFP. Notably, we observed an increase in spike-phase coupling in the higher frequencies (40-100 Hz) in the active versus passive condition in neurons that were strongly sound modulated (see methods), consistent with a behavioral state modulation of spike timing (**Fig. 4A-B**, two-sided Wilcoxon signed-rank test, FDR corrected *p <* 0.05). Surprisingly, even non-sound modulated cells showed enhanced coupling albeit at a narrower range of gamma frequencies and with a lower coupling strength (**Fig. 4B**). We further explored how spatial location modulated spike phase-coupling and found a significant increase in the coupling between gamma frequency activity and spiking activity when the AM sound came from the rewarded location specifically (**Fig. 4F**; two-sided Wilcoxon signed-rank test, FDR corrected *p <* 0.05), and that this effect was absent in the passive block. These results suggest that there is a spatially selective engagement of neuronal assemblies in the auditory cortex that is also state dependent. Next, we assessed how gamma changed over cortical depth using a mixed effect model (see methods). This model included location (rewarded vs non-rewarded), cortical depth, behavioral condition, and their interactions. Spike–gamma coupling was significantly increased during active engagement (*β* = 11.353, SE = 4.007, *t* = 2.834, *p* = 0.0048) and was strongest for sounds presented at the rewarded location (*β* = 26.543, SE = 8.898, *t* = 2.983, *p* = 0.0030). Coupling decreased with cortical depth overall (*β* = -3.489, SE = 1.396, *t* = -2.500, *p* = 0.0128), and this effect was modulated by reward location, as reflected in a significant location × depth interaction (*β* = -3.575, SE = 1.800, *t* = -1.986, *p* = 0.0477). Rewarded location further interacted with behavioral condition (*β* = -13.397, SE = 5.677, *t* = -2.360, *p* = 0.0187). In contrast, the depth × condition interaction was not significant (*β* = 0.999, SE = 0.814, *t* = 1.228, *p* = 0.220), and the three-way interaction among location, depth, and condition showed a trend-level effect (*β* = 2.067, SE = 1.160, *t* = 1.782, *p* = 0.0755). Given the theta-gamma coupling findings, we next investigated whether theta frequency phase also contributes to spike coupling (**Fig. S8C**).

**Figure 4.**
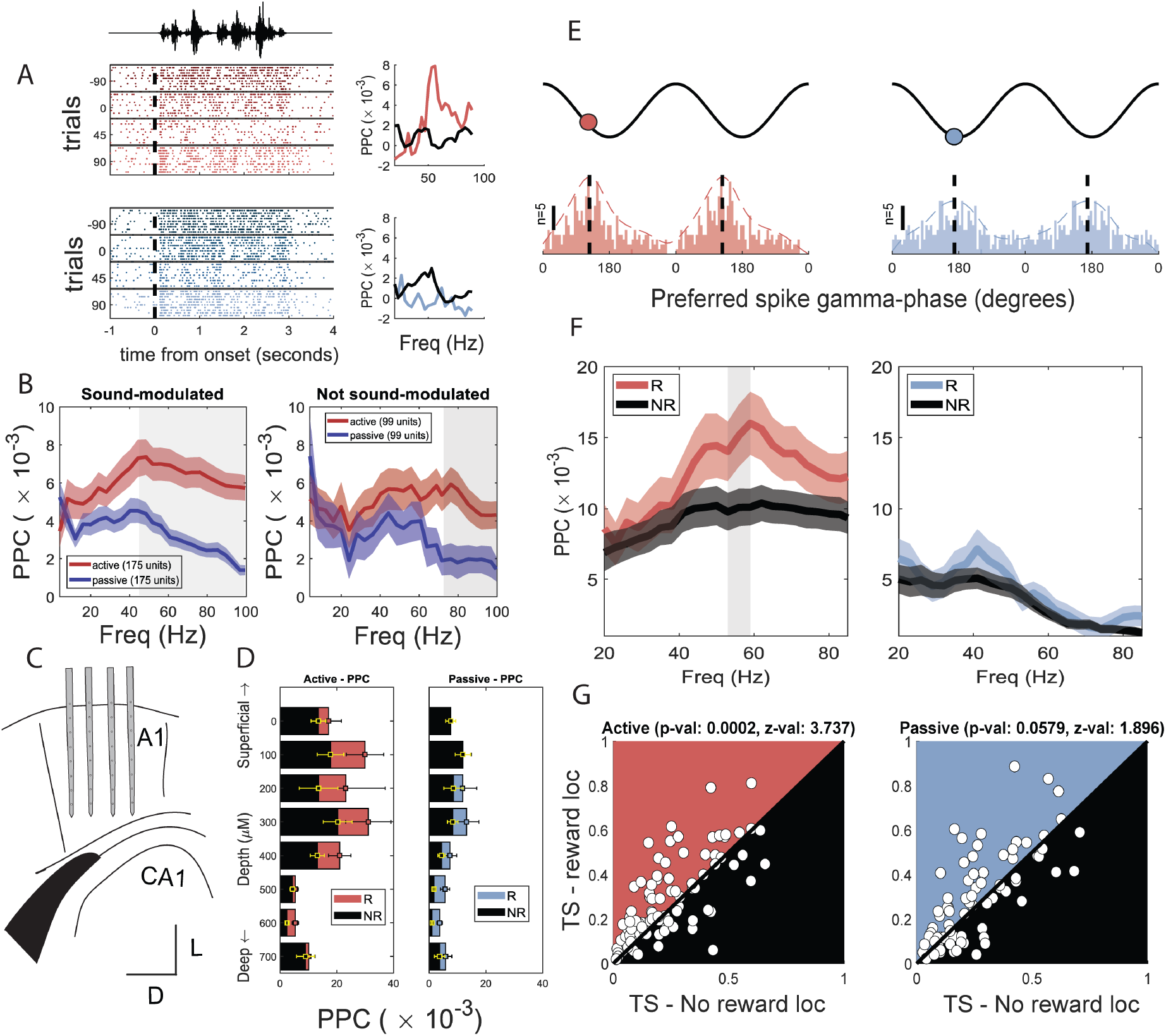
Neurons selectively couple to gamma for stimuli delivered from salient spatial locations. **A**. Example neuron that shows enhanced coupling between the active block and passive block that is spatially dependent. Pairwise phase consistency (PPC) values for rewarded and not rewarded locations are shown to the right. **B**. PPC coupling for sound modulated (*left*) and non-sound modulated neurons (*right*) in the active versus passive conditions. **C**. Cartoon illustration showing location of shanks implanted in the auditory cortex. **D**. Spike-gamma coupling across depths in the active (*left*) and passive conditions (*right*). PPC contributed from the rewarded location is shown in red while all other locations are noted in black. **E**. Preferred phase of gamma for all units in active and passive condition when sound was presented in the 90 degrees location. Note a shift to an earlier phase in the gamma cycle that is associated with greater temporal alignment. **F**. PPC spectrum for rewarded versus not-rewarded location in active (*left*) and passive (*right*) task conditions (shaded area at 53-59 Hz: *p <* 0.05, Wilcoxon sign-rank test) **G**. Trial similarity differences across locations for active (*left*) and passive blocks (*right*), colored triangles indicate 90 degree rewarded location whereas black triangle indicates all other non-rewarded locations. We found no differences in the distribution around the unity line for passive state but a robust increase in similarity for the rewarded location relative to non-rewarded locations.

We found that spikes coupled to theta selectively for the behaviorally relevant location during the active block, with no significant differences observed in the passive block. This result may reflect the nesting of gamma within theta oscillations that has been previously associated with attentional engagement^37^. Together, these results indicate that behavioral engagement in our task is associated with theta-gamma coupling and elevated spike-gamma coupling in superficial cortical layers that encode spatial saliency.

We next assessed whether this enhanced coupling has functional consequences for sensory encoding. To do so, we measured the reliability of the response of strongly phased-coupled neurons to AM sounds. To this end, we applied the measure of trial similarity^36^. Specifically, we randomly divided the trials in each location into two equal groups, binned spike times with a time resolution of 25 milliseconds, and calculated Pearson’s correlation coefficients between the two resulting peristimulus time histograms (PSTHs). This process was repeated 100 times to obtain a mean correlation coefficient, or trial similarity. Our results reveal that neurons with stronger gamma coupling exhibited increased trial-to-trial reliability, as measured by correlations between independently sampled spike trains. This increase in reliability was specific to the rewarded location (*p* = 0.0002, *z* = 3.7) in the task-engaged condition but was absent in the passive condition (**Fig. 4G**; *p* = 0.0579; *z* = 1.896), indicating an increase in reliability of the response that is dependent on behavioral engagement. These results indicate that gamma synchronization organizes spike timing to reduce trial-by-trial variability and enhance the fidelity of sensory representation encoding.

### Temporal coordination, rather than firing rate, mediates spatial selectivity

To better understand the behavior of individual neuron responses, we examined how the firing rate of neurons were modulated by behavioral state and location. Firing rate changes have been shown to be modulated by relevant stimuli in the auditory cortex and can be expressed as both increases and decreases in firing rate^7,45^. We captured the extent of modulation by condition using a Contextual Modulation Index (CMI, see methods), where positive values indicate a facilitation, whereas negative CMI values denote suppression of the neuronal response in the active block. Despite robust modulation of temporal dynamics, we did not observe significant differences in the modulation of firing rate by location across the entirety of the neural population (**Fig. 5A,5F**) Instead, individual neurons showed heterogeneous changes, with increases and decreases balancing at the population level. We observed a balanced percentage of neurons in the population that increased and decreased firing rates for each location (**Fig. S5B**). More-over, we divided neurons into narrow waveform (NW) and broad waveform (BW) (**Fig. S4B-D**) and found that the direction of the CMI could not be explained by putative cell identity (**Fig. S4E**).

**Figure 5.**
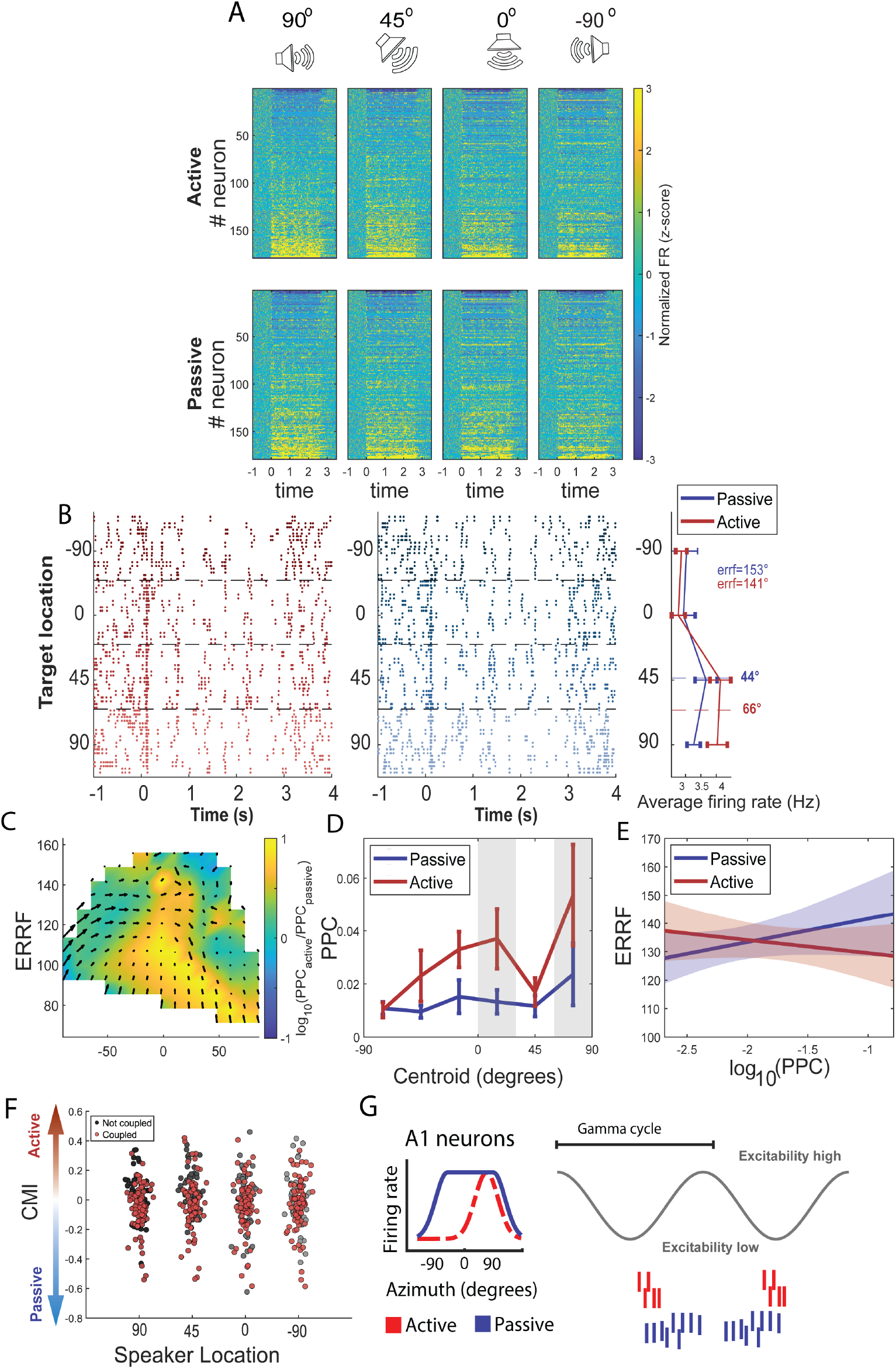
Gamma-coupling is selective to neurons representing the rewarded spatial location. **A**. Average firing rate across locations for active (*top*) and passive (*bottom*) conditions for all units recorded sorted from most inhibited to most activated. **B**. Raster plots from an example cell showing AM sound evoked sound responses for each location in passive (*left*) and active (*right*) conditions. **C**. Vector field map indicating trajectory of centroid and ERRF from passive to active. Heatmap indicates change of PPC from passive to active and the relationship between PPC coupling strength, ERRFs and centroid. Specifically, contralateral tuned neurons that are significantly coupled to gamma did not change their centroids and have very narrow receptive fields, whereas ipsilateral tuned neurons do not couple to gamma and move their centroids in the contralateral direction, while broadening their receptive fields. **D**. Average PPC values for each centroid bin, neurons whose centroid was closer to the rewarded location show an increase in spike-phase coupling to gamma frequencies. Shaded areas denote significant differences between active and passive: *p <* 0.05 corrected by FDR). **E**. Interaction plot showing the relationship between ERRF and the strength of PPC coupling at gamma frequencies. Shaded areas show confidence intervals for adjusted ERRF. ERRF was lower during the active relative to the passive block (*β* = -25.03, *p* = 0.033), with a significant condition × PPC interaction (*β* = -12.89, *p* = 0.028). **F**. Average CMI value across conditions for all neurons sorted by the 4 speaker conditions in the active condition **G**. Working model that reflects the main findings of the study. We found neurons tightly coupled to gamma during periods of high cortical excitability narrow their receptive fields during task-engagement and are tuned to the behaviorally relevant location.

To further investigate how task conditions influence spatial tuning, we examined the relationship between spiking rate and spatial receptive field properties. Each neuron was characterized using previously published tuning metrics including: centroid (i.e., the location at which neurons fire the most) and equivalent rectangular receptive field (ERRF, a measure of spatial breadth of receptivity)^45,75,82^(**Fig. 5B, Fig. S6-7, S8B**). We reasoned that cells that are more or less weakly tuned to a specific location, might undergo rate modulation for those locations specifically. Therefore, we separated cells by their spatial tuning preference and re-analyzed the populations using this criterion and still found no significant effect of rate-location interactions as measured by CMI (**Fig. S8D**). These results provide evidence that gamma oscillation modulation associated with behavioral engagement in our task design does not specifically modify spatial tuning sensitivity via a rate coding strategy.

Given that we found strong changes in LFP-spiking interactions that were modulated by location, we analyzed how temporal coordination influenced spatial tuning by examining the relationship between spike–field coupling and receptive field properties. We specifically looked at the relationship between tuning features and gamma spike-field coupling. We observed that neurons with stronger gamma coupling (PPC*>*0.002, **Fig. S8A**) exhibited narrower receptive fields and tuning centered near the rewarded location (**Fig. 5C-D**). Moreover, our results revealed significant effects of condition and spike-field coupling on ERRF (**Fig. 5E**). Relative to the passive block, ERRF was significantly lower during the active block (*β* = -25.032, SE = 11.554, *t* = -2.167, *p* = 0.033). PPC alone did not show a significant main effect on ERRF (*β* = 8.242, SE = 5.539, *t* = 1.488, *p* = 0.140). However, there was a significant interaction between condition and PPC (*β* = -12.886, SE = 5.759, *t* = -2.237, *p* = 0.028), indicating that the relationship between spike-field coupling and receptive field size differed across behavioral states. Specifically, PPC was positively related to ERRF in the passive condition, whereas this relationship was reversed in the active condition. These findings suggest that spatial saliency can reshape spatial auditory representations through enhanced temporal precision.

## DISCUSSION

Our results demonstrate that task-engaged spatial selectivity in A1 is expressed primarily through enhanced temporal coordination rather than robust changes in firing-rate encoding by individual neurons. Behaviorally relevant locations selectively recruited neural populations into gamma-synchronized activity without a corresponding and consistent increase in firing rate. Notably enhanced spike-field coupling increased response reliability selectively for rewarded spatial locations, directly linking oscillatory dynamics to the fidelity of sensory representations. Spatial modulation of spike-field coupling was most prominent in superficial cortical layers, and coupling strength was systematically related to both neuronal reliability and spatial tuning properties. Together, these findings provide strong support for temporal coordination as a mechanism through which the auditory cortex selectively prioritizes encoding of behaviorally relevant spatial information.

Despite compelling evidence that A1 employs temporal coding strategies in sound processing, direct demonstration that temporal coding serves as a critical carrier of behaviorally relevant spatial information has not been shown. While the importance of precise timing and coordinated neuronal activity for discriminating competing sound sources has been shown^59^, previous studies have not attempted to disentangle the relative contribution of firing-rate and temporal coding components that support spatial representations during behavior. Here, we directly dissociate these relative contributions. We observed heterogeneous changes in firing rate across the neural population, with approximately half of neurons increasing and half decreasing their activity during task engagement. This is consistent with prior work revealing that auditory task engagement produces co-existing excitation and suppression effects at the population level during auditory task engagement^38,45,81^. Our findings, combined with this work, effectively rule out a generalized rate-based account of selective spatial processing. Instead, we found support for enhanced temporal coordination that was selectively aligned to behaviorally relevant locations and predicted both response reliability and spatial tuning. Unlike visual cortex, where attentional states are often reflected through firing-rate gain^22,72,73^, our findings indicate that behaviorally relevant locations are preferentially represented through enhanced temporal coordination in A1, establishing spike timing rather than firing-rate gain as the dominant coding dimension underlying spatial prioritization during behavior.

This distinction may be particularly important for the auditory cortex because auditory signals unfold dynamically over time and do not occupy stable spatial coordinates like tonotopy. Temporal coordination could therefore provide a more flexible coding strategy by selectively enhancing the timing relationships among neurons representing behaviorally relevant inputs while preserving the distributed architecture necessary for complex auditory scene analysis. Our findings further demonstrate that gamma oscillations play a central role in mediating this temporal coordination. Gamma-band activity arises through interactions between excitatory and inhibitory interneurons, particularly parvalbumin-positive (PV) cells^14,16^. Beyond reflecting local processing, gamma synchronization has been implicated in attention, perceptual selection, and sensory gain control across modalities^9,73^. In the auditory cortex, rhythmic entrainment of neural activity to sound structure and attentional rhythms further supports the idea that temporal alignment is a key mechanism for selective processing^43,71^. Gamma synchronization may thus function as a temporal binding signal that transiently links neurons encoding behaviorally relevant spatial locations into coherent perceptual representations. Such transient synchronization would allow for distributed neuronal populations to dynamically coordinate their activity without requiring a fixed anatomical spatial map. Crucially, gamma bursts emerge across coupled cortical circuits with matched timing and phase relationships, and instantaneous phase relationships between regions can selectively gate information flow^62^. Gamma synchronization may therefore provide a dynamic routing mechanism through which distributed cortical populations transiently organize into functionally coherent ensembles selective for behaviorally relevant locations.

Consistent with this framework, neurons exhibiting stronger gamma coupling also show increased response reliability, suggesting that temporal coordination improves the signal-to-noise ratio of auditory representations by reducing trial-to-trial variability and stabilizing sensory encoding^6,78^. Temporal synchronization may therefore not simply reflect heightened excitability, but instead actively organizes how information is represented and routed across cortical populations. Such coordination may be particularly important in speech perception and auditory scene analysis, where behaviorally relevant information unfolds across multiple timescales spanning phonetic, syllabic, and prosodic structure^33^. Dynamic synchronization could allow the auditory system to selectively enhance relevant speech streams while preserving sensitivity to competing sources in complex listening environments.

Our laminar analyses further revealed that spatial modulation of spike-field coupling was strongest in superficial layers. These layers are known to integrate corticocortical and top-down inputs, suggesting that behaviorally relevant signals may influence auditory processing through feedback pathways^8,13,51^. This interpretation is also consistent with predictive coding frameworks in which higher-order cortical areas dynamically shape sensory representations in early sensory cortex through top-down feedback signals^19,29^. Within this architecture, enhanced gamma synchronization in superficial cortical layers may reflect the selective stabilization of behaviorally relevant sensory representations. Previous work suggests that gamma-band coordination in superficial layers can preferentially route temporally coherent activity while suppressing less coordinated outputs^3^, providing a potential mechanism through which distributed cortical populations dynamically prioritize behaviorally relevant representations over competing inputs. In our task, consistent association between reward and a specific spatial location likely generated a priority signal that biased auditory cortical processing toward that region of space. Importantly, these effects generalized across neutral stimuli, indicating that modulation operated at the level of spatial representation rather than stimulus-specific encoding. As such, our findings are best interpreted within a framework of value-driven saliency, in which behavioral relevance shapes sensory processing independently of stimulus identity^4,25^. This interpretation is further supported by changes in pupil diameter that tracked behavioral state^50,64^. Increased spike-gamma coupling was associated with elevated arousal, consistent with studies showing that neuromodulatory systems, particularly the LC–NE system, regulate cortical gain and network dynamics^5,38,50,77^. Elevated arousal may facilitate the selective synchronization of neural populations representing behaviorally relevant stimuli.

We also observed selective enhancement of theta-gamma coupling for behaviorally relevant spatial locations during task engagement. This effect may be interpreted in at least two, not mutually exclusive ways. First, it is consistent with evidence from the visual attention literature showing that frontal theta oscillations modulate gamma activity in sensory cortex to co-ordinate rhythmic sampling and attentional selection, with direct consequences for behavioral performance^26,27,35^. The enhanced theta-gamma coupling observed here may serve a similar role in the auditory domain, whereby top-down theta signals dynamically coordinate sensory processing at behaviorally relevant spatial locations consistent with findings from other attentional tasks^37^. Second, theta oscillations occur on a timescale closely aligned with the temporal structure of speech, particularly its syllabic rhythm^33,48^. Theta-gamma coupling may therefore support the parsing of complex auditory information by aligning slower speech-related temporal structure with faster gamma-band activity encoding finer-grained sensory representations. Together, our findings suggest that enhanced theta-gamma coupling during task engagement may reflect the combined influence of attentional coordination and temporally structured sensory encoding.

These findings have direct implications for auditory scene analysis and the biological underpinnings of the cocktail party problem, where listeners must selectively extract relevant sound sources from competing complex acoustic inputs^10,11,49,59^. Auditory cortical representations are strongly influenced by behavioral relevance and sound category, indicating that spatial hearing is not purely stimulus-driven but also depends on top-down factors that guide listening behavior^21,40^. Task-engagement can rapidly reshape receptive fields, sharpening spatial tuning and enhancing encoding of task-relevant features in A1^30–32,45^. Spatial selectivity in A1 is also not an intrinsic neuronal property but is itself behaviorally contingent, with neurons exhibiting mixed selectivity and context-dependent spatial tuning^17,47^. One potential solution allowing such elasticity is that behaviorally relevant spatial information is represented not through stable anatomical organization, but through the dynamic coordination of distributed neuronal populations. Temporally coordinated ensembles can emerge transiently and flexibly, allowing distributed populations of neurons to selectively group and route behaviorally relevant information without requiring stable spatial segregation. Selective synchronization of spike timing could therefore organize neurons representing behaviorally relevant locations into functional ensembles while preserving the distributed architecture necessary for flexible auditory scene analysis. Our results framed within this context suggest spatial selectivity may be achieved at least in part not through global elevation of firing rates, but through selective temporal coordination of distributed neuronal populations representing behaviorally relevant spatial locations.

While we used distinct behavioral contexts to isolate the impact of task-engagement, the relative contribution of selective attention versus general changes in arousal state remain difficult to disentangle. Notably, alpha–beta power differed across behavioral states but not across locations, suggesting that generalized task engagement alone is unlikely to account for the location-specific changes in gamma synchronization and spike–field coupling reported here. Future studies will therefore be necessary to isolate the relevant contributions of these factors to selective processing of sound location. In addition, although task engagement enhanced response precision and reliability, we did not obtain a direct behavioral readout of spatial discrimination performance. Establishing this relationship will be important for determining the behavioral significance of the temporal coordination mechanisms identified here. An important open question is whether the task-dependent gamma synchronization we observe is generated locally within A1 or inherited through top-down projections from higher-order auditory or prefrontal regions. Future work combining cell-type-specific manipulations and causal circuit approaches will be necessary to resolve the details of this mechanism.

In summary, our findings suggest that temporal coordination provides a solution to a fundamental computational constraint of auditory spatial processing. Because A1 lacks a stable topographic representation of space, rate coding could inherently prove to be a relatively inefficient strategy. Without a spatial scaffold to index location, spike counts would be less efficiently read out subsequently reducing spatial information^54,58^. Gamma-mediated synchronization offers an alternative strategy by transiently organizing distributed neuronal populations into functionally coherent ensembles selective for behaviorally relevant locations. More broadly, these findings suggest that dynamic temporal coordination may represent a general principle through which cortical systems flexibly prioritize relevant information when sensory representations are distributed rather than topographically organized.

## METHODS

### Subjects

All procedures involving animals were approved by the University of Illinois at Urbana-Champaign Institutional Animal Care and Use Committee (IACUC). A total of 17 transgenic mice were used in this study. Original breeding pairs of parvalbumin-Cre (PV-Cre: B6;129P2-Pvalbtm1(cre)Arbr/J), somatostatin-Cre (SST-Cre: STOCK Ssttm2.1(cre)Zjh/J), and all breeding was carried out within our facility. The experimental cohort included both male and female progeny consisting of SST-Arch (n = 8), SST-Cre (n = 4), and PV-Cre (n = 5) mice. At the time of the recording sessions, all animals were between 12 and 16 weeks old. These animals were selected so they could be used in subsequent optogenetic studies but no optogenetic manipulations were performed in these experiments.

### Intrinsic signal imaging

Under 1.5-2% isoflurane anesthesia, stereotaxic surgery was performed to install a custom head-plate and intrinsic signal imaging was carried out following approaches described in earlier studies^57^, employing a custom tandem lens macroscope (composed of Nikkor 55 mm 1:2.8 and 85 mm 1:2) and a 16-bit CMOS monochrome camera (BFLY-U3-23S6M-C, FLIR). Before imaging, mice were given a subcutaneous injection of chlorprothixene at a dosage of 1.5 mg/kg body weight, and isoflurane was reduced to 0.75%. A custom well was created using Body Double “Fast Set” silicone and a glass coverslip to maintain a saline pool and prevent evaporation. The camera was first focused on the vasculature between the skull and the surface of the auditory cortex and then focused to a depth 400*µ*m below the cortical surface. The intrinsic signals were captured at a frequency of 20Hz under red light illumination (625nm). Each trial consisted of a 1.5 s baseline, followed by a 1 s pure-tone stimulus (75 dB SPL at 3, 10, or 30 kHz), and a 28.5 s inter-trial interval. Acoustic stimuli were delivered via a Tucker Davis Technologies (TDT; Alachua, FL) RZ6 processor and TDT MF1 Multi-Field Magnetic Speakers operated in closed-field mode. Images captured during the response period (0.5s to 2.5s from the sound onset) were averaged and divided by the average image during the baseline to generate frequency response maps. These maps were then binarized and color-coded to construct tonotopic gradients using custom MATLAB scripts.

### Electrode implantation surgery

Following intrinsic imaging, stereotaxic surgery was performed to implant the electrode array. The permanent head-plate was mounted 3mm anterior to bregma and secured to the skull with three stainless steel screws and dental acrylic. A fourth screw connected to a metal pin was implanted in the skull above the contralateral cerebellum to serve as a reference. We delineated the boundary of the primary auditory cortex (A1, right hemisphere) with intrinsic signal imaging. A 32-contact electrode array from Neuronexus (model: a 4 × 8-5mm-100-400-177-CM32), featuring 100*µ*m inter-contact spacing and 400 *µ*m shank spacing, was precisely inserted into A1 perpendicular to the cortical surface using a stereotaxic arm. Because of the curvature of the auditory cortex surface, inserting all four shanks to the same depth was not always feasible. The probes were advanced until all electrode contacts were embedded within the cortical tissue. After surgery, the mice were given a recovery period of 4-7 days before being habituated to head fixation, as described below.

### Auditory Task Training

Mice were gradually restricted to a single hour of water access over a 7-day period following headplate habituation. Training was conducted in a sound booth, where animals were presented with rewarded trials playing 0.5 s pure tones at 10,035 Hz and non-rewarded trials playing human speech–enveloped white noise, delivered at 5-second intertrial intervals. For rewarded trials, the animal had up to 2 seconds after sound onset to operantly lick the water port and trigger a reward release; responses occurring later did not result in reward delivery. Each training session included 162 pure-tone presentations, allowing for a maximum of 162 possible rewards. Water rewards consisted of 10 *µ*L droplets of 0.02% saccharin solution. Mice were trained under these conditions until they responded to more than 80% of tone trials for three consecutive days. This training phase typically lasted 7–10 days. Once criterion was met, animals underwent a sham recording session to habituate them to the procedures and events associated with recording days.

### Auditory stimuli

Auditory stimuli consisted of three stimuli: two discrete neutral 3 second long amplitude modulated broadband white noise stimuli convolved with different human speech envelopes and one rewarded a 0.5 s pure tone at 10,035 Hz. All stimuli were generated in MATLAB and delivered through four TDT ES-1 electrostatic speakers. The non-rewarded AM sounds were shaped using temporal envelopes extracted from human speech recordings in the Harvard IEEE speech corpus, a resource commonly used in cocktail party effect studies^1^. Stimulus intensity was calibrated prior to each session with a conditioning amplifier and microphone (Brüel & Kjær, Nærum, Denmark; amplifier model 2690, microphone model 4939-A-011), ensuring a sound level of 75 dB SPL at the mouse’s head. Playback duration was 3 s for the non-rewarded white noise stimuli and 0.5 s for the pure tone, that applied a 1 ms cosine ramp applied at onset and offset.

### Recording and Data Acquisition

Electrophysiology recordings were acquired using the TDT RZ10X system within a soundproof chamber. Broadband neural activity from all 32 channels was sampled at 24,414.0625 Hz. Each recording session consisted of 162 rewarded pure tone trials and 80 non-rewarded trials presenting 40 human speech–enveloped white noise stimuli for each distinct envelope. During the recording, the four speakers were equidistantly placed 18 cm from the mouse’s head and arranged along the azimuthal plane: one directly in front (0°), two contralateral (45° and 90°), and one ipsilateral (-90°) relative to the right auditory cortex recording site. Pure tone stimuli were presented from a single fixed location for the entire recording day, whereas non-rewarded white noise stimuli were delivered randomly across all four speakers, with playback balanced such that each speaker was used 10 times for each envelope by the end of the session. Each non-rewarded trial consisted of 3 s of stimulus playback followed by 2 s of silence, while each rewarded trial consisted of 0.5 s of stimulus playback followed by 4.5 s of silence. Pupil recordings were obtained using a FLIR Blackfly S BFS-U3-16S2C camera controlled by SpinView software. Videos were acquired at 20 Hz and triggered by the TDT RZ10X system to synchronize with other experimental events.

### Spike Sorting

Spike sorting was performed with Kilosort 2.5 to automatically detect multi-units^60^. The resulting clusters were then manually curated in Phy to determine whether they represented genuine neural activity or noise^67^. Clusters showing licking-related artifacts or uniform activity across channels were classified as noise, and artifact-like waveforms were carefully excluded from clusters that otherwise contained neural signals. Clusters recorded on the same channel that overlapped in principal component feature space and whose merged cross-correlogram exhibited a refractory period were merged.

The curated cluster data were subsequently imported into MATLAB from Phy using the spikes toolbox, with each cluster assigned to the channel where its average spike amplitude was highest. To evaluate unit quality, the sorting Quality toolbox was used to identify single-units (SUs) based on isolation distance and L-ratio^70^. Units were classified as SUs if they exhibited fewer than 5% of inter-spike intervals shorter than 2 ms, an isolation distance greater than 15, and an L-ratio below 0.25. In cases where isolation distance and L-ratio were not available, the inter-spike interval criterion alone was applied. These classification standards are consistent with prior single-unit activity studies (^42,56^. Clusters not meeting these criteria were categorized as multi-units (MUs). Neurons were deemed as sound-modulated if either their average or max firing rate during sound playback was significantly different (t-test, p-value below 0.05) from that of baseline.

### Pupil Tracking

Pupillometry videos were collected and analyzed using the MATLAB-based pupillometry toolbox described by^66^. Briefly, pupil contours were detected frame-by-frame and fit with a circular model using the MATLAB procedure to estimate pupil center and radius. This fitting approach enabled robust extraction of pupil diameter across time, even in the presence of partial occlusion or noise, providing a continuous measure of pupil dynamics for subsequent analyses.

### Local Field Potentials preprocessing

Data acquired was low pass filtered at 500 Hz and notch filtered at 60 Hz, then resampled to 1024 Hz. Highly noisy electrodes were rejected after visual inspection. Trials contaminated with more than 1 lick within sound playback were rejected as well as those trials whose RMS was higher than 5 standard deviations from the mean. We calculated the power spectral density of LFP signals for each channel in A1. We fit the PSD using the FOOOF algorithm between 1 and 150 Hz to isolate the oscillatory peaks above a 1/f background^23^.

### Gamma-event detection

Discrete gamma event detection was performed using the CBASS tool developed by^65^. Multi-channel LFP was filtered in gamma band (40-60 Hz) and candidate events were selected at the troughs of the filtered signal in a reference channel. The spectrotemporal dynamics underlying each candidate event was parameterized using the real and imaginary part of the analytical representation (Hilbert function in MATLAB) of the filtered signal in each channel. An optimization procedure is then applied to find the threshold yielding the most significant distance between events having a low and high enrichment score (fraction of time that events fall into an enriched partition). Events above threshold were retained

### Spike-Field Coupling

The phase locking of spikes to LFP features at each frequency was measured for individual units using the phase from 4 to 100 Hz in bins of 4 Hz at the time of each spike^79^. Only neurons that fired at least 50 spikes during stimulus presentation were used. Reference LFP was taken by the two electrodes 200 um away from the electrode where the unit was recorded to minimize spike energy leakage into the LFP. The mean angle of the phases for a given neuron’s spikes was taken as the preferred phase. We considered neurons to be significantly coupled if their PPC was higher than 0.002, similar to previous studies^52^.

### Contextual modulation index

We quantified contextual modulation for each speaker location L using the measure Contextual Modulation Index (CMI) defined as:

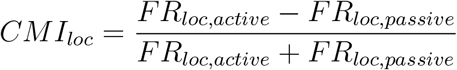

Where FR is the response to the AM sound coming from speaker location L in either the active condition or passive condition. This measure varies from -1 complete suppression to +1 maximal facilitation and has a value of 0 when the condition has no effect on the response. Firing rates were by definition always non-negative.

### Spatial tuning features

The spatial tuning curve of each auditory neuron was obtained by averaging firing rates recorded during playback of the AM sound at each speaker location. To quantify the spatial tuning properties of each neuron, two metrics were used: centroid, and equivalent rectangular receptive field (ERRF). These metrics have been widely used in previous studies of auditory spatial processing^45,82^.

Spatial tuning was further characterized by the centroid. The centroid was calculated as follows: the peak-response range was identified as the set of speaker locations that elicited firing rates within 75% of the unit’s maximum response. These locations were represented as vectors weighted by their corresponding firing rates. The vector sum of these weighted responses was then computed, and the direction of the resultant vector was defined as the centroid. The breadth of spatial tuning was quantified by the width of the equivalent rectangular receptive field (ERRF), defined as the width of a rectangle with an area equal to that under the spatial tuning curve and a height equal to the peak firing rate.

To estimate confidence intervals for centroid and ERRF width spike rates at each speaker location were resampled non-parametrically with replacement 500 times, preserving the number of trials at each location. Each parameter was computed for every resampled dataset, yielding a distribution of 500 estimates per metric. The mean of each distribution was taken as the final parameter value, and the 2.5th and 97.5th percentiles were used to define the 95% confidence intervals.

### Phase-Amplitude Coupling

We quantified phase-amplitude coupling (PAC) for all channels using the modulation index^18,74^(MI). For each channel, we extracted the LFP starting at -1000 until 4000 ms following the presentation of the sound. We filtered each trial within the respective frequency bands of interest (using pop eegfiltnew.m from EEGLAB, v.2019.1). We extracted the instantaneous phase from the low-frequency signal and the amplitude from the high-frequency component using the Hilbert transform. Trials were cropped to the final window of interest (0 to 3000 ms relative to the sound onset), ensuring that edge artifacts are removed. Next, we concatenated the phase and the amplitude signals across trials and computed the MI as described previously (18 phase bins). We computed MI separately for each location and condition. To standardize MI in each channel, frequency, and conditions, we computed 200 surrogate MIs by randomly combining the phase and amplitude from different trials (trials-shuffling), separately for different locations. These values were then used to produce z transforms of raw MI values. Standardizing MI values eliminates potential systematic differences that might be reflected in PAC across locations.

## Statistical Analysis

All statistical analyses were performed using MATLAB R2025a. The significance threshold was set at 0.05 throughout. Descriptive statistics are reported as mean ± standard deviation unless otherwise noted. Comparisons of lick rates across sound source locations were assessed using a one-way ANOVA (Fig. 2B). Differences in pre-stimulus spectral power, CMI, and trial similarity between active and passive blocks were assessed using a two-sided Wilcoxon signed-rank test (Fig. 2D; Fig. 3F; Fig. 4G). LFP spectral differences between active and passive conditions were evaluated using a cluster-based permutation tests in which condition labels were randomly exchanged across 5000 permutations. The effects of spatial location on LFP gamma power and theta gamma phase-amplitude coupling were assessed using linear mixed-effects models (LME) with location as a fixed effect and subject as a random effect; model coefficients are reported in the main text and figure descriptions. Effects of location and condition on pupil diameter were assessed using a two-way ANOVA with subject included as a random factor (Fig. 2G-H). Differences in PPC between conditions and locations were assessed using two-sided Wilcoxon signed-rank tests with FDR correction for multiple comparisons across frequency. The influence of behavioral state, spatial location, cortical depth, and their interactions on spike-gamma coupling was modeled using a linear mixed-effects model with subject as a random effect (Fig. 4D). Spatial receptive field properties were compared across behavioral conditions and spike-field coupling levels using a linear mixed-effects model, including condition, PPC and their interaction as fixed effects and subject as a random effect (Fig. 5E).

## ACKNOWLEDGMENTS

This work was funded by National Science Foundation grants: SMA-2319321 (H.J.G and K.S), and EFRI-2515342 (H.G.). M.I. was funded by the National Science Foundation (2123781), the Beckman Institute Graduate Fellows Program, and the Arnold and Mabel Beckman Foundation.

## AUTHOR CONTRIBUTIONS

M.I, Z.Q., KS, and H.J.G. designed the experiments. M.I, Z.Q. and H.J.G. collected the data. M.I. analyzed data. M.I. and Z.Q. curated data. K.S., S.S., and H.J.G. acquired funding. M.I., Z.Q. and H.J.G. wrote the manuscript. K.S. and S.S. provided revisions to the manuscript.

## DECLARATION OF GENERATIVE AI AND AI-ASSISTED TECH-NOLOGIES

During the preparation of this work the author(s) used Claude and ChatGPT in order to improve readability. After using this tool/service, the author(s) reviewed and edited the content as needed and take(s) full responsibility for the content of the published article.

## DATA AND CODE AVAILABILITY

All data and code supporting the findings from this study are available upon reasonable request from the corresponding author (H.J.G).

