## Supplemental Information for "Representations of spatial saliency in auditory cortex are selectively organized through temporal coordination"

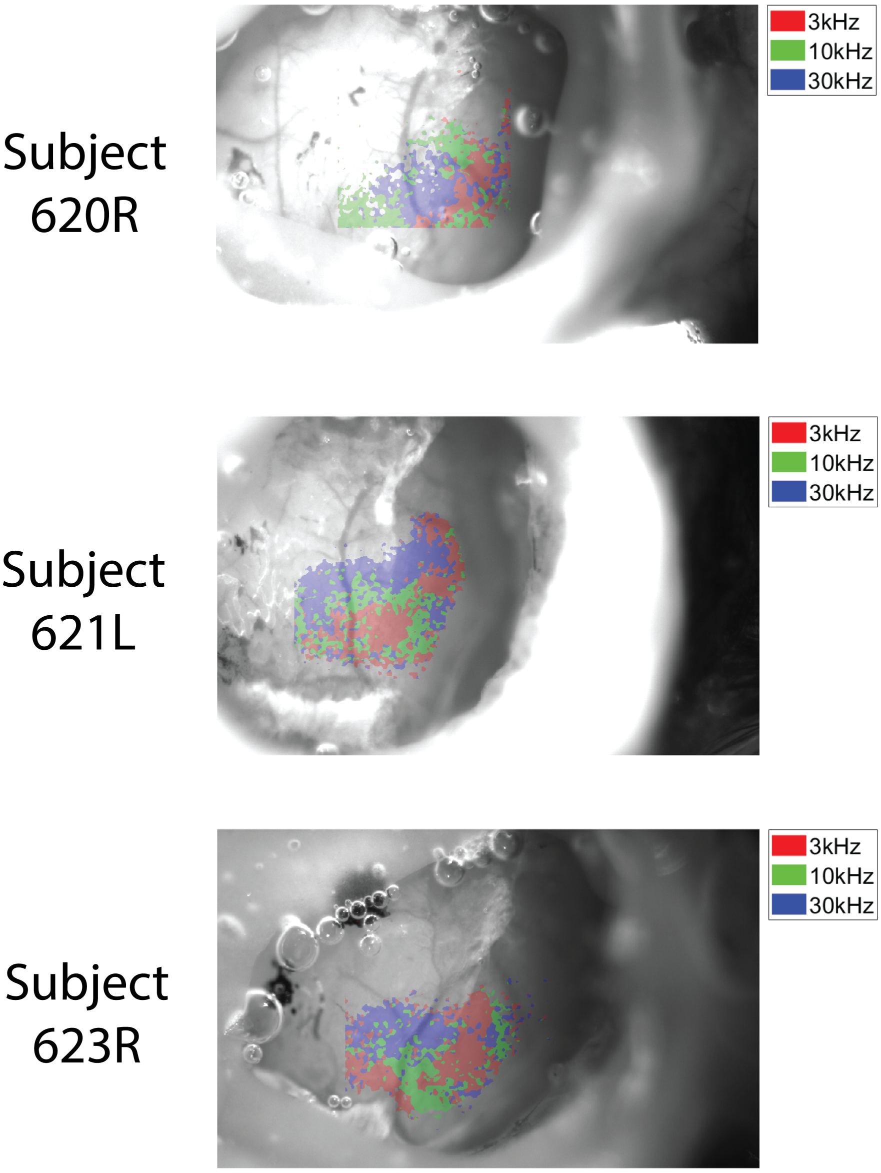


**Figure S1. Intrinsic imaging from representative mice used in this study**. Thinned skull intrinsic signal images collected from A1 from three example mice (subjects **620R, 621L and 623R**). Pseudo-color masks represent hemodynamic response to auditory stimuli delivered to the contralateral ear (see methods). Color boundaries delineate areas that are maximally activated by one of three different tone frequencies delivered: (red: 3 kHz; green: 10 kHz; blue 20 kHz).

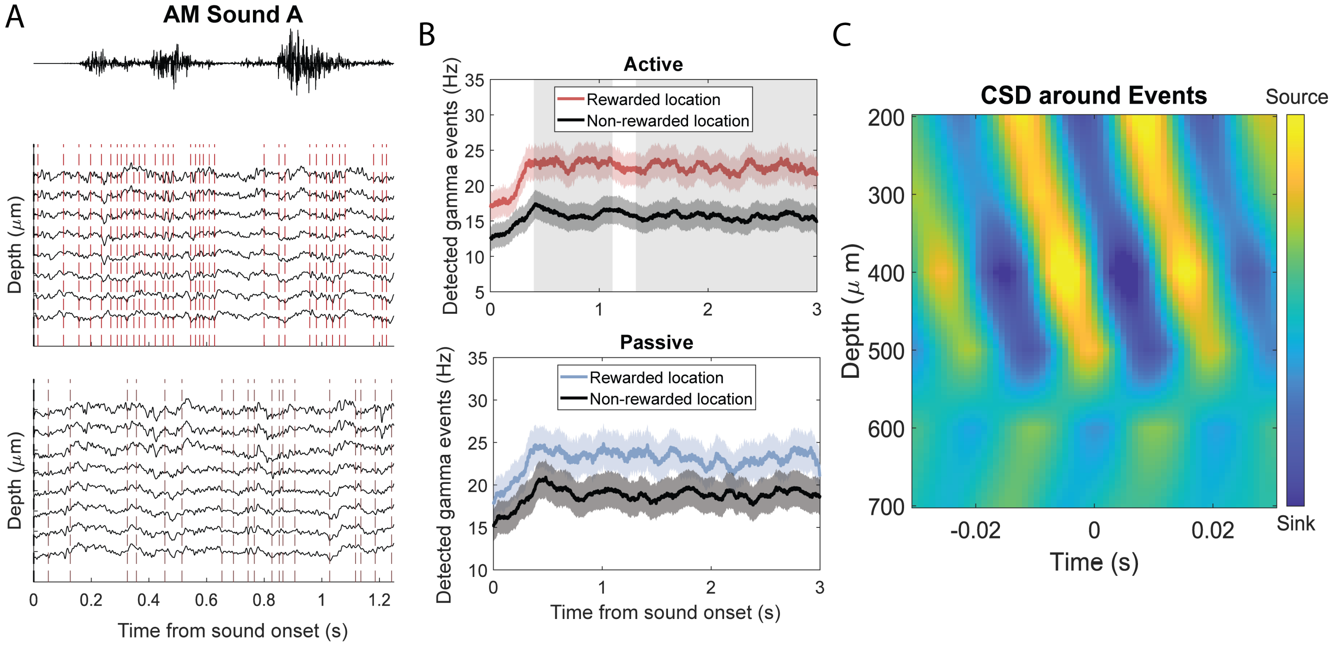


**Figure S2. Gamma event detection. A.** Example electrophysiology voltage traces from a representative mouse used in this study aligned to the AM sound A stimulus. Multichannel LFP events filtered in the gamma (40-60 Hz) range were used for event detection. CBASS analysis was used to determine "gamma events". Analysis was applied to recordings collected from A1 during the presentation of a sound from the rewarded (*top*) and non-rewarded (*bottom*) location during the task-engaged active block. Events were selected at the troughs of the filtered signal relative to a reference channel and are denoted by dashed lines. Note the overall increase in "events" for the rewarded location compared to non-rewarded locations. **B**. CBASS-detected gamma event rate during sound presentation for active and passive blocks. We observed an increase in event rates that remained elevated during the sound presentation window (≈3 seconds) in both active and passive blocks although only the active block showed spatial significance. Data from (n=17) mice. Shaded areas denote significant differences ($p < 0.01$, cluster-based permutation test) between rewarded and non-rewarded locations. **C**. CSD depth profile from a representative animal time-locked to detected gamma events using CBASS. We found that detected events were primarily localized to the channels spanning granular to superficial layers.

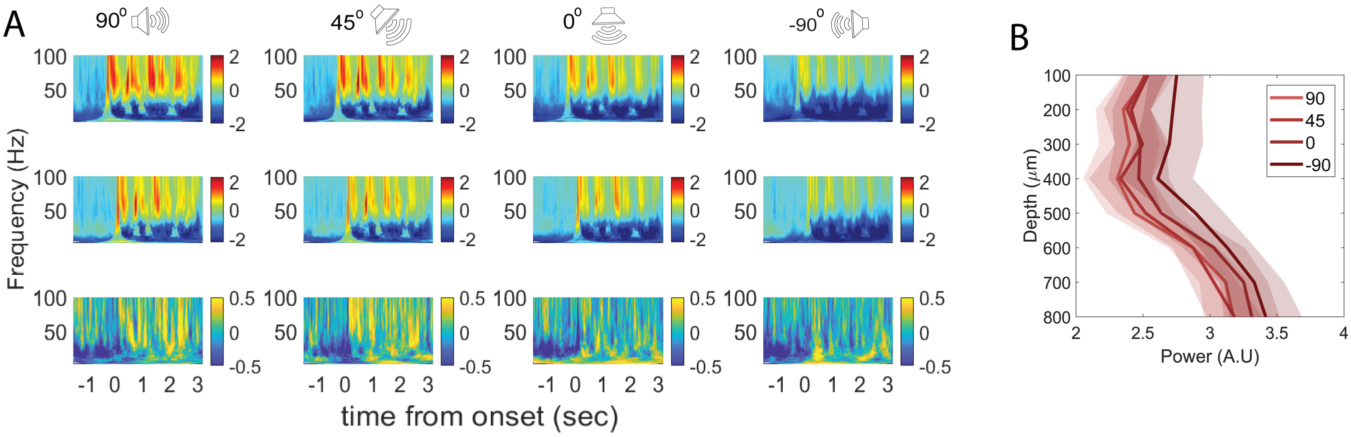


**Figure S3**. LFP power across active and passive states by location. **A.** Normalized time-frequency spectrograms for AM sound A in active (*top*) and passive (*middle*) blocks. Difference plots between active and passive states are shown below (*bottom*) for each spatial location. Warm colors (yellow) on the bottom row represent frequencies where the active condition shows higher relative power than the passive condition. Note a reduction in warm colors as the stimulus is played further away from the 90-degree rewarded location. **B.** Active period theta (4-9 Hz) power across the depth profile for the 4 sound source locations.

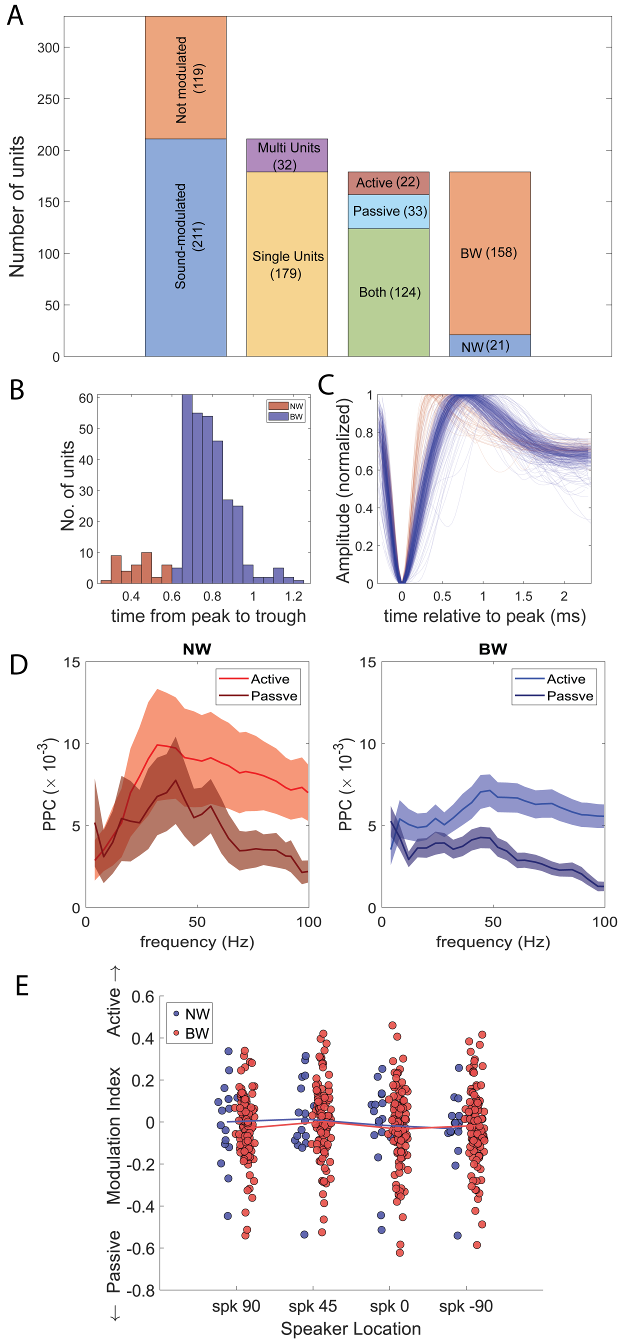


**Figure S4.** Cell classification and waveform analysis for all cells recording in this study. A. Categories of recorded cell types: (n=330) cells from a total of (n=17) subjects. Cells were quantified in terms of the number of neurons in each category including: sound modulated, single units, behavioral state modulated, and waveform (narrow: **NW** or broad: **BW**) structure. **B**. Action potential (AP) waveform classification. Histogram demonstrating distribution of peak-to-trough temporal profiles for all single-unit cells. Neurons were classified based on timing of peak to trough responses as narrow waveform (NW) or broad-waveform (BW) units. **C**. Average AP waveforms for all neurons classified as BW (blue) and NW (red) collected in this study. **D**. PPC analysis sorted by cell waveform type. Across active and passive conditions, there was significantly increased coupling by NW (*left*) units compared to BW (*right*) units. Both waveform types also showed substantially elevated coupling in active versus passive sessions. **E**. Modulation index for NW and BW neurons across the four spatial locations. We did not find significant changes in modulation index for either waveform type.

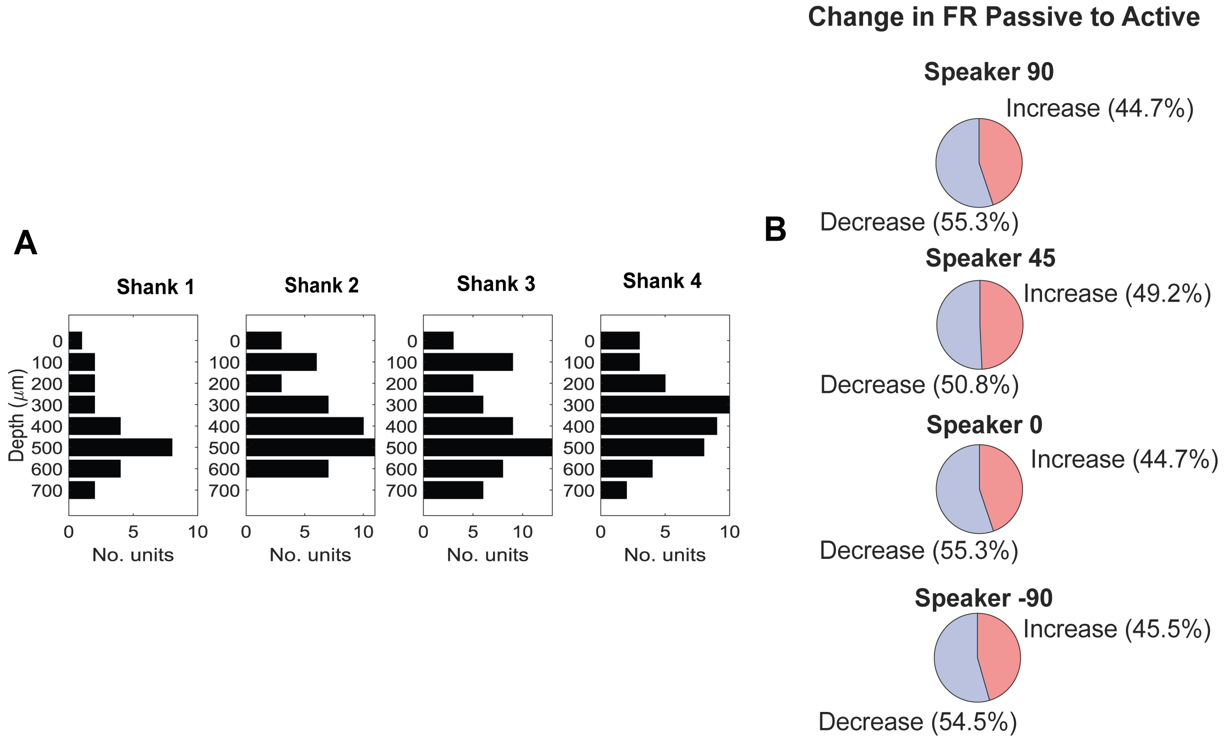


**Figure S5. Cell densities across cortical depth. A.** Number of recorded units identified across the four shanks sorted by depth from the cortical surface. We collected a broad distribution of cells across all layers. **B.** Proportion of units that showed significant PPC coupling (PPC threshold of $> 0.002$) during the active period and their relative change in firing rate between active and passive sessions. We found that neurons were fairly balanced across the population with slightly more units showing a slight reduction in stimulus evoked firing rate compared to the passive state on average.

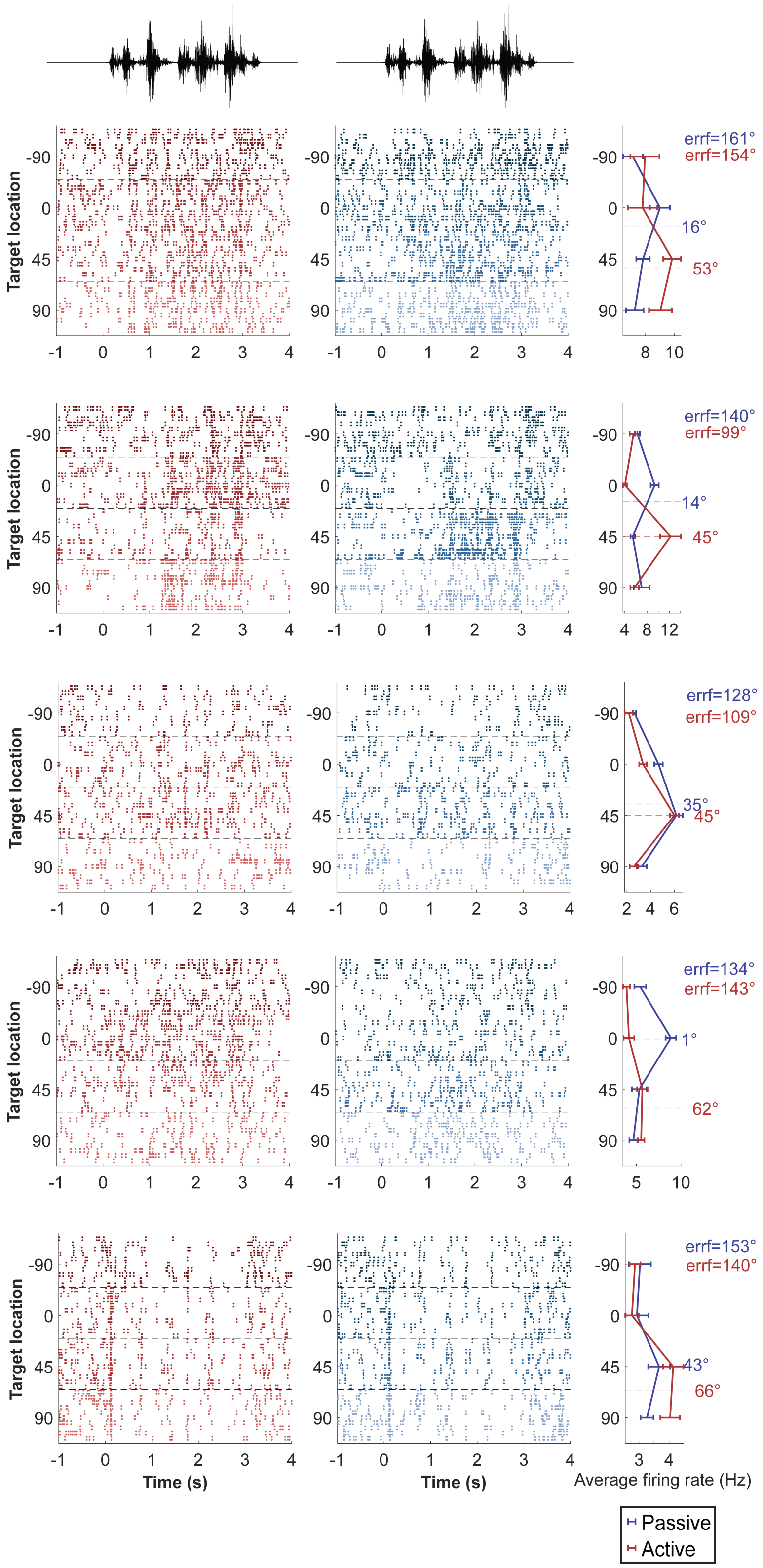


**Figure S6. Representative example neurons recorded from A1 in active and passive sessions for <u>AM sound A.</u>** All units shown demonstrated elevated spike-phase coupling in the active versus passive blocks (PPC$> 0.002$). Raster plots during active (red) and passive (blue) sessions for each speaker location. For each sound, the corresponding spatial tuning firing rate for each of the 4 speaker locations is shown to the right of the raster plots. The ERRF and centroid were calculated for each representative neuron and are shown to the right. Most neurons show a narrowing of ERRF and a shift in centroid towards the 90 degree location.

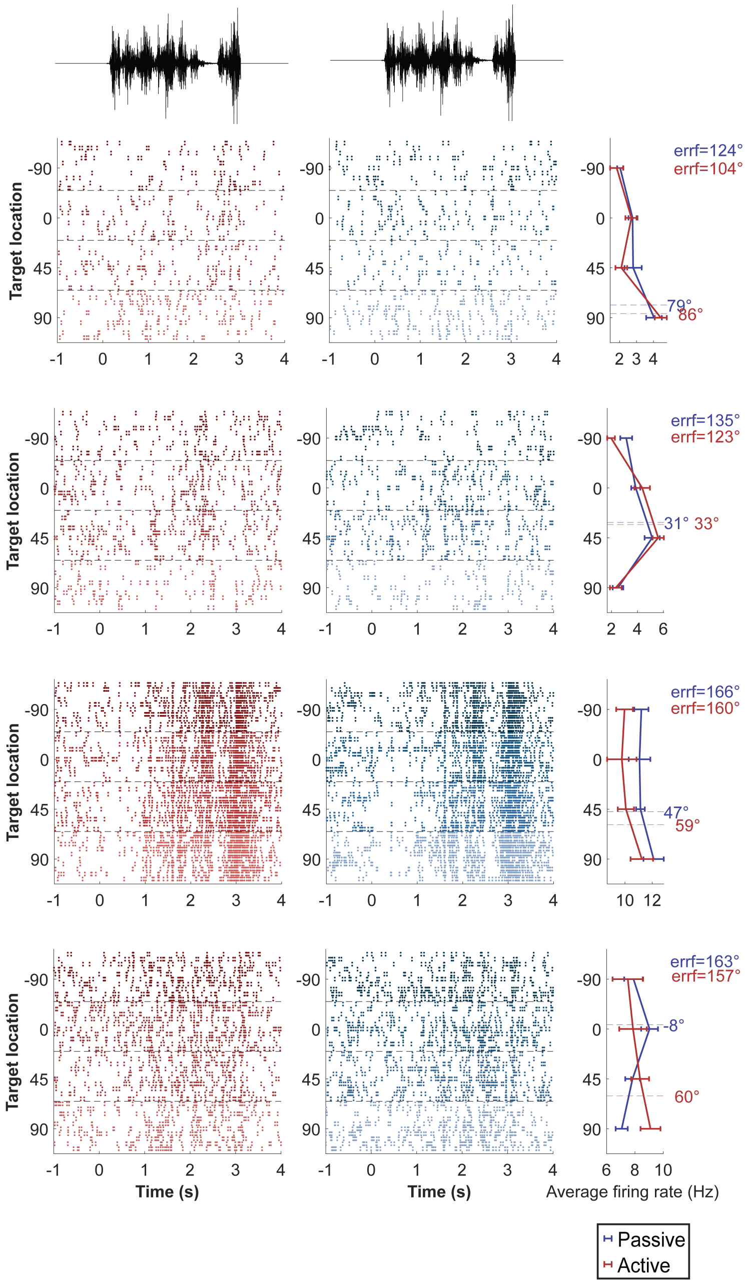


**Figure S7. Representative example neurons recorded from A1 in active and passive sessions for <u>AM sound B</u>.** Cells selected demonstrated elevated spike-phase coupling in the active versus passive blocks (PPC$> 0.002$). Raster plots during active (red) and passive (blue) sessions for each speaker location. For each sound, the corresponding spatial tuning firing rate is shown to the right of the raster plots for the 4 speaker locations. The ERRF and centroid were calculated for each representative neuron. Most PPC elevated neurons show a narrowing of ERRF and a shift in centroid towards the 90 degree location.

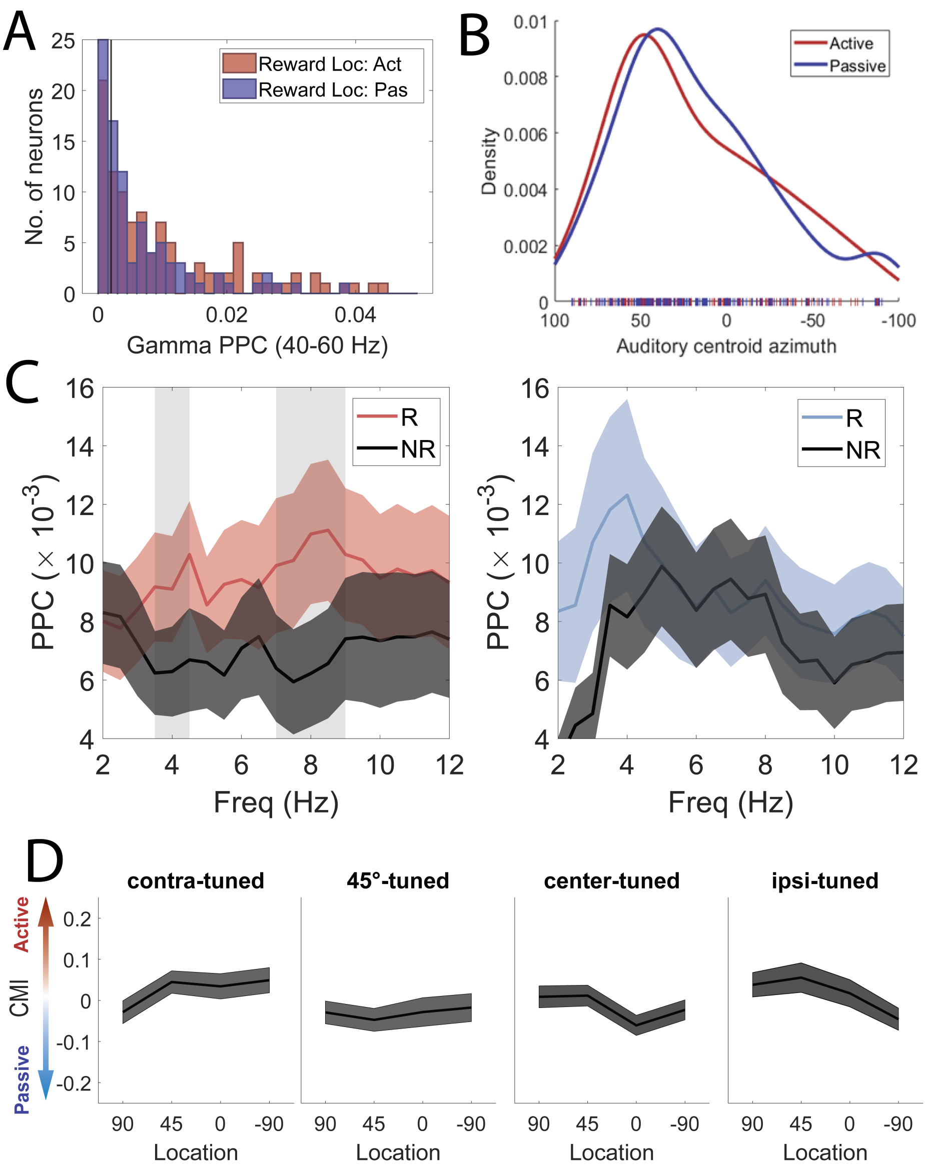


**Figure S8. Phase coupling strength and relative spatial tuning of recorded units. A.** Distribution of gamma PPC for all units during the active and passive sessions. Note a shift to a higher coupling strength in the active period. A PPC threshold of $> 0.002$ was used to identify significantly coupled units for subsequent analyses. **B.** Density plot showing the centroid distribution of neurons in passive (blue) or active (red) sessions. Note that the centroids for this population shift closer to the rewarded location (90 degrees) as animals transition from passive to active sessions. Tickmarks represent the centroid for each unit in passive (blue) or active (red) conditions. **C.** PPC values at lower frequencies (2-12hz) for active (*left*) and passive (*right*) sessions for rewarded (red or blue) and non-rewarded (black) locations. Shading reflects frequency range of significantly elevated coupling between rewarded and non-rewarded locations (two-sided Wilcoxon sign rank test, $p < 0.05$ FDR-corrected). Note increases are also found for delta and theta ranges consistent with the AM modulation components primarily inherent in speech and found in the spectral envelopes of each AM stimulus. **D.** Contextual modulation index (CMI) at each location for neurons sorted by their spatial tuning preference. Gray bars represent standard error from the mean. We observed no significant effect of location on firing rate.
